## Supplementary Table 1 for "Generation and Trapping of a Mesoderm Biased State of Human Pluripotency"

### Table 1 Single Cell qPCR assay list

| Assay ID | Gene Symbol | RefSeq | Amplicon Length | Detects gDNA | Best Coverage |
|---|---|---|---|---|---|
| **Hs01060665_g1** | *ACTB* | NM_001101.3 | 63 | Yes | Yes |
| **Hs00154192_m1** | *BMP2* | NM_001200.2 | 60 | No | Yes |
| **Hs03676628_s1** | *BMP4* | NM_130850.2;NM_001202.3; NM_130851.2 | 116 | Yes | Yes |
| **Hs01034913_g1** | *BMPR1A* | NM_004329.2 | 94 | Yes | Yes |
| **Hs00193796_m1** | *CER1* | NM_005454.2 | 92 | No | Yes |
| **Hs01897804_s1** | *CITED2* | NM_001168388.2;NM_001168389.2;NM_006079.4 | 106 | Yes | Yes |
| **Hs00607528_s1** | *CLDN6* | NM_021195.4 | 154 | Yes | Yes |
| **Hs00164004_m1** | *COL1A1* | NM_000088.3 | 66 | No | Yes |
| **Hs00976734_m1** | *CXCR4* | NM_003467.2;NM_001008540.1 | 153 | No | No |
| **Hs00171876_m1** | *DNMT3B* | NM_001207055.1;NM_001207056.1;NM_175848.1;NM_175849.1;NM_175850.2;NM_006892.3 | 55 | No | Yes |
| **Hs00172872_m1** | *EOMES* | NM_001278183.1;NM_001278182.1;NM_005442.3 | 81 | No | Yes |
| **Hs01549976_m1** | *FN1* | NM_212482.1;NM_054034.2; NM_002026.2;NM_212478.1; NM_212474.1;NM_212476.1 | 81 | No | Yes |
| **Hs00232764_m1** | *FOXA2* | NM_021784.4;NM_153675.2 | 66 | No | Yes |
| **Hs00255287_s1** | *FOXD3* | NM_012183.2 | 78 | Yes | Yes |
| **Hs00173503_m1** | *FRZB* | NM_001463.3 | 108 | No | Yes |
| **Hs00246256_m1** | *FST* | NM_006350.3;NM_013409.2 | 108 | No | No |
| **Hs00544355_m1** | *GAL* | NM_015973.3 | 125 | No | Yes |
| **Hs00171403_m1** | *GATA4* | NM_002052.3 | 68 | No | Yes |
| **Hs00232018_m1** | *GATA6* | NM_005257.4 | 91 | No | Yes |
| **Hs00906630_g1** | *GSC* | NM_173849.2 | 100 | No | No |
| **Hs00193435_m1** | *HAS2* | NM_005328.2 | 63 | No | Yes |
| **Hs00242160_m1** | *HHEX* | NM_002729.4 | 110 | Yes | Yes |
| **Hs00705137_s1** | *IFITM1* | NM_003641.3 | 93 | Yes | Yes |
| **Hs01547673_m1** | *ITGA5* | NM_002205.2 | 54 | No | Yes |
| **Hs00761767_s1** | *KRT19* | NM_002276.4 | 116 | Yes | Yes |
| **Hs00764128_s1** | *LEFTY1* | NM_020997.3 | 136 | Yes | Yes |
| **Hs00745761_s1** | *LEFTY2* | NM_001172425.1;NM_003240.3 | 102 | Yes | Yes |
| **Hs00355202_m1** | *LGALS1* | NM_002305.3 | 63 | No | Yes |
| **Hs00232144_m1** | *LHX1* | NM_005568.3 | 60 | No | Yes |
| **Hs00702808_s1** | *LIN28A* | NM_024674.4 | 143 | Yes | Yes |
| **Hs00430824_g1** | *MIXL1* | NM_031944.1 | 152 | No | No |
| **Hs00899658_m1** | *MMP1* | NM_001145938.1;NM_002421.3 | 64 | No | Yes |

| Assay ID | Gene Symbol | RefSeq | Amplicon Length | Detects gDNA | Best Coverage |
|---|---|---|---|---|---|
| **Hs01548727_m1** | *MMP2* | NM_004530.4;NM_001127891.1 | 65 | No | Yes |
| **Hs01085598_g1** | *MYL7* | NM_021223.2 | 74 | No | Yes |
| **Hs04399610_g1** | *NANOG* | NM_024865.2 | 101 | Yes | No |
| **Hs00378379_m1** | *NCLN* | NM_020170.3 | 65 | No | Yes |
| **Hs00415443_m1** | *NODAL* | NM_018055.4 | 68 | No | Yes |
| **Hs00219496_m1** | *PAF1* | NM_019088.3;NM_001256826.1 | 100 | No | Yes |
| **Hs04260367_gH** | *POU5F1* | NM_001173531.1;NM_002701.4;NM_203289.4 | 77 | Yes | Yes |
| **Hs01375212_g1** | *RPS18* | NM_022551.2 | 93 | Yes | Yes |
| **Hs00183425_m1** | *SMAD2* | NM_001135937.2;NM_001003652.3;NM_005901.5 | 129 | No | No |
| **Hs00195591_m1** | *SNAI1* | NM_005985.3 | 66 | Yes | Yes |
| **Hs00751752_s1** | *SOX17* | NM_022454.3 | 149 | Yes | Yes |
| **Hs01053049_s1** | *SOX2* | NM_003106.3 | 91 | Yes | Yes |
| **Hs00610080_m1** | *T* | NM_001270484.1;NM_003181.3 | 132 | No | Yes |
| **Hs00761239_s1** | *TAGLN2* | NM_001277224.1;NM_001277223.1;NM_003564.2 | 163 | No | Yes |
| **Hs02339499_g1** | *TDGF1* | NM_003212.3;NM_001174136.1 | 170 | Yes | No |
| **Hs00902257_m1** | *WNT3* | NM_030753.4 | 76 | No | Yes |
