## Supplementary Fig for "Generation and Trapping of a Mesoderm Biased State of Human Pluripotency"

### Supplementary Figure Legends

**Figure 1: Single Cell qPCR analysis.**

Heatmap analysis of 232 single cells analysed across 45 genes using a Fluidigm BioMark system, hierarchical clustering was performed for the genes assessed. The heatmap visualises the individual gene expression after normalisation across genes and samples. White coloured genes indicates undetected levels. Black arrows indicate cells which appear to express pluripotency associated genes (particularly *SOX2, NANOG* and *POU5F1*) at a high level whilst also expressing early differentiation markers. Cells co-expressing pluripotency and differentiation associated genes were readily detected in the *MIXL1*(**+**)/SSEA-3(**+**) fraction but not the other fractions.

**Figure 2: Assessing the stem cell potential of *MIXL1*(+)/SSEA-3(+) substate.**

**a)** Live TRA-1-81 staining fluorescent images of colonies derived after the first passage into a 48 well plate at 4x, TRA-1-81(RED) and *MIXL1*-GFP (GREEN). Wells marked with white stars indicates clones that survived the passage and stained positive for TRA-1-81. Of the 44 colonies passaged, 27 survived and stained positive for TRA-1-81. **b)** Immunofluorescent analysis of NANOG expression in HES3 *MIXL1*-GFP clones 2-D2 and 3-C6 growing in E8V conditions. Merged images display Hoechst (Nuclei) in blue and NANOG positive cells in red. Secondary only staining control is also shown. **c)** Bar chart showing the percentage positive cells for the stem cell associated antigens BF4, CD9, SSEA-3, SSEA4, TRA-1-60s, TRA-1-81 and TRA-2-49 for six clonal lines established (Mean of all lines +/- SD). All lines displayed high expression of these surface markers.

**Figure 3: Averaged qPCR Signature Comparison**

The average 1/Ct values for 47 genes from single cell qPCR analysis. Genes were ordered from highest to lowest expression based on the *MIXL1*(-)/SSEA-3(+) fraction. A solid line connects the mean expression points to give a state “signature” with surrounding shaded area represents the 95% confidence interval of the data. **a)** Displays the state signatures of *MIXL1*(-)/SSEA-3(+)(red), *MIXL1*(+)/SSEA-3(+) (green) and *MIXL1*(+)/SSEA-3(-) (blue) grown in MEF/KOSR conditions. **b)** Displays the state signatures of *MIXL1*(+)/SSEA-3(+) cells grown in MEF/KOSR (green) and Primo (purple) conditions. The state signature of the both *MIXL1*(+)/SSEA-3(+) were very similar.

**Figure 4: Single Cell Gene Expression Plots.**

The single cell gene expression distribution was similar between the two *MIXL1*(+)/SSEA-3(+) fractions from MEF/KOSR (green) and Primo (purple) conditions. 1/Ct values for each single cell for a given gene. Mean and standard deviation are displayed on top of data sets as black bars. Cells are split into their respective sorted fractions MEF/KOSR conditions *MIXL1*(-)/SSEA-3(+) cells in red, *MIXL1*(+)/SSEA-3(+) cells in green, *MIXL1*(-)/SSEA-3(+) cells in blue and Primo conditions *MIXL1*(+)/SSEA-3(+) cells in purple. **a)** Contains a collection of plots from genes associated with pluripotency. **b)** Contains a collection of plots from key genes associated with mesendoderm differentiation. **c-f)** Contains plots from the remaining genes assessed by single cell qPCR.

**Figure 5: Differentiation Time Course**

Flow cytometry density plots of HES3 *MIXL1*-GFP cells stained for SSEA-3 at indicated time points after induction of differentiation in E8 containing 3µM CHIRON. Red boxes indicate the sorting gates for each timepoint. The expression of *MIXL1*-GFP increases first, before the eventual loss of SSEA-3.

**Figure 6: Clones generated from the *MIXL1*(+)/SSEA-3(+) from Primo medium exhibit normal stem cell growth and characteristics.**

**a )** Live TRA-1-81 staining fluorescent images of colonies derived from single cell deposition of *MIXL1*(+)/SSEA-3(+) from PRIMO Plus conditions after the first passage into a 48 well plate at 4x, TRA-1-81(RED) and *MIXL1*-GFP (GREEN). Wells marked with white stars indicates clones that survived the passage and stained positive for TRA-1-81. **b)** Flow cytometry density plot of *MIXL1*-GFP versus SSEA-3 from clone 12-F11 grown in MEF/KOSR conditions. **c)** Bar chart of percentage positive stem cell associated antigen SSEA-3 and *MIXL1*-GFP expression for five clonal lines during initial expansion in MEF/KOSR conditions. **d)** Flow cytometry density plot of *MIXL1*-GFP versus SSEA-3 from clone 12-F11 after being transitioned into E8V conditions. **e)** Bar chart of percentage positive stem cell associated antigens BF4, CD9, SSEA-3, SSEA4, THY-1 and TRA-1-81 for six clonal lines. All lines displayed high expression of these surface markers.

**Figure 7: Components of Primo can be substituted for others that target the same pathway.**

**a-e)** Flow cytometry density plots of T-venus and SSEA-3 expression in different conditions. **a)** In standard E8V conditions. **b)** Using PRIMO Plus formulation, IWP2 was replaced for DKK1 at 100ng/mL, density plot reveals high double expression 4 days after the first passage. **c)** CHIR99021 was replaced with SB216763 at 10μM, density plot reveals high double expression after 3 days of induction. **d)** LPA was replaced with S1P, density plots demonstrate the ability of S1P to block differentiation, 1.92μM S1P maintained a high proportion of double positive cells after 3 days of induction (optimal concentrated highlighted with a red box). **e)** LPA was replaced with GRI977143, density plots demonstrate the ability of GRI to block differentiation, 4μM GRI maintained a high proportion of double positive cells after 3 days of induction (optimal concentrated highlighted with a red box).

**Figure 8: Passage 10 NANOG and SOX2 expression Analysis**

Cells in all conditions at the tenth passage show high expression of NANOG and SOX2. **a-b)** Immunofluorescence analysis of Hoechst, *MIXL1*-GFP, and **a)** NANOG or **b)** SOX2 expression of HES3 *MIXL1*-GFP cells in PRIMO Plus, E8+LPA (0.96μM), E8 alone, E8 with 1μM IWP-2 added and E8+LPA (0.96μM) (Secondary antibody only staining) for 3 days post to 9 passages in PRIMO Plus. A merged image of all three channels is present below

Hoechst (Blue), *MIXL1*-GFP (Green) and NANOG or SOX2 (Red). **c-d)** Stacked percentage bar charts displaying cell profiler analysis of 3 wells for each condition (Bars are mean ± SD, n= 3 technical repeats) for *MIXL1*-GFP, and **c)** NANOG or **d)** SOX2 expression grown in PRIMO Plus, E8+LPA (0.96μM), E8 alone and E8 with 1μM IWP-2 added, for 3 days post to 9 passages in PRIMO Plus.

S1

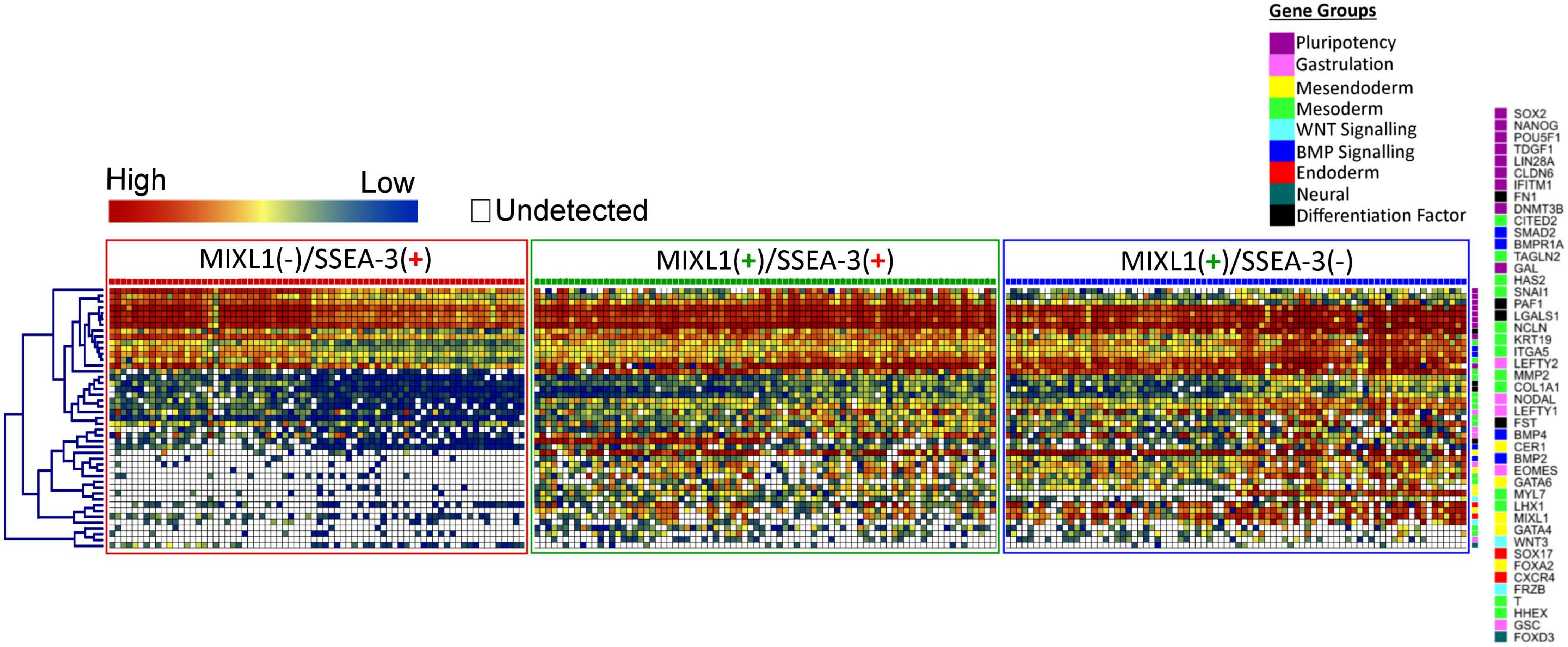
High
Low
Undetected
Gene Groups
Pluripotency
Gastrulation
Mesendoderm
Mesoderm
WNT Signalling
BMP Signalling
Endoderm
Neural
Differentiation Factor
MIXL1(-)/SSEA-3(+)
MIXL1(+)/SSEA-3(+)
MIXL1(+)/SSEA-3(-)
SOX2
NANOG
POU5F1
TDGF1
LIN28A
CLDN6
IFITM1
FN1
DNMT3B
CITED2
SMAD2
BMPR1A
TAGLN2
GAL
HAS2
SNAI1
PAF1
LGALS1
NCLN
KRT19
ITGA5
LEFTY2
MMP2
COL1A1
NODAL
LEFTY1
FST
BMP4
CER1
BMP2
EOMES
GATA6
MYL7
LHX1
MIXL1
GATA4
WNT3
SOX17
FOXA2
CXCR4
FRZB
T
HHEX
GSC
FOXD3

S2

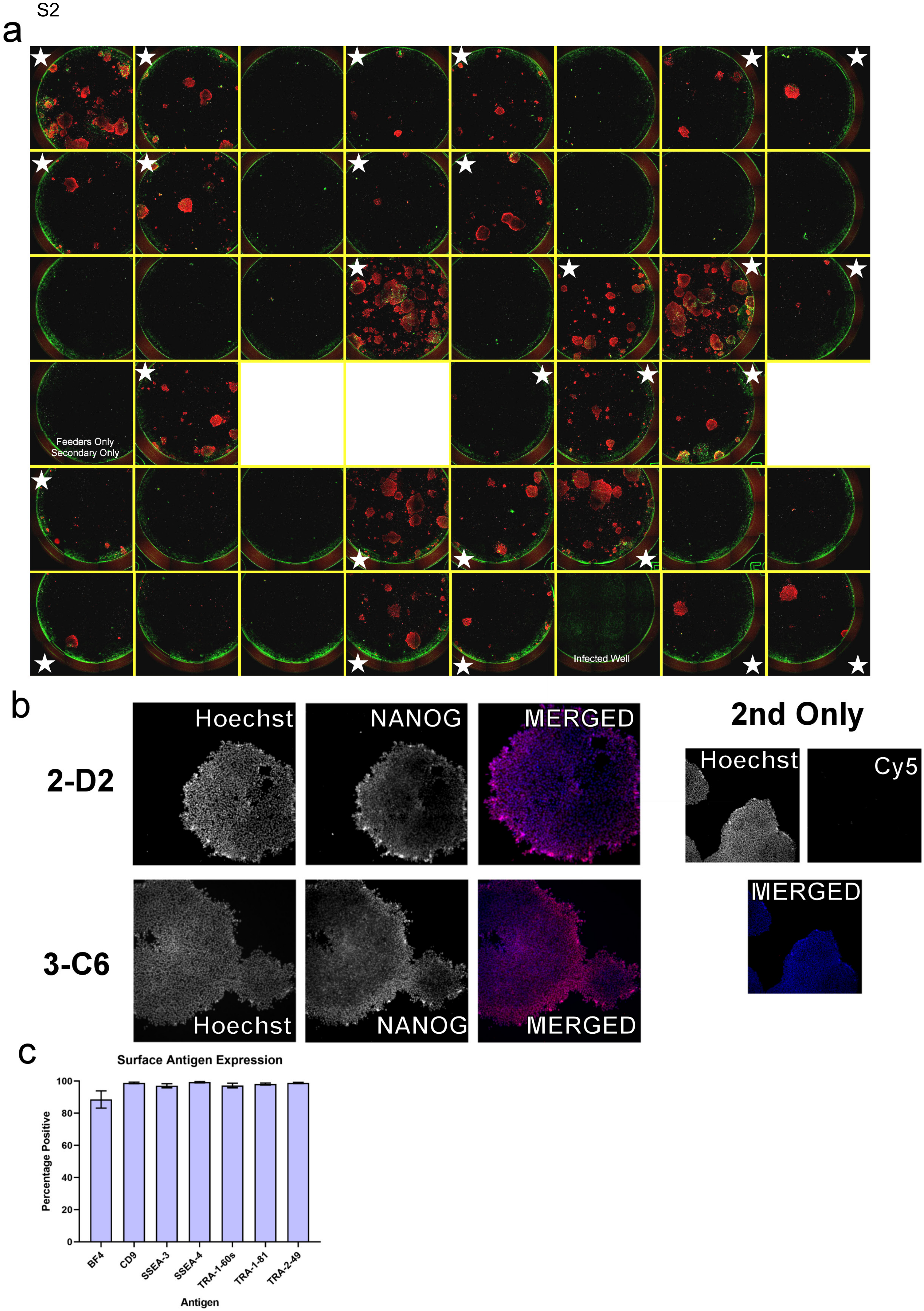
a
Feeders Only
Secondary Only
Infected Well
b
2-D2
3-C6
Hoechst
NANOG
MERGED
2nd Only
Hoechst
Cy5
MERGED
c
Surface Antigen Expression
Percentage Positive
0
20
40
60
80
100
BF4
CD9
SSEA-3
SSEA-4
TRA-1-60s
TRA-1-81
TRA-2-49
Antigen

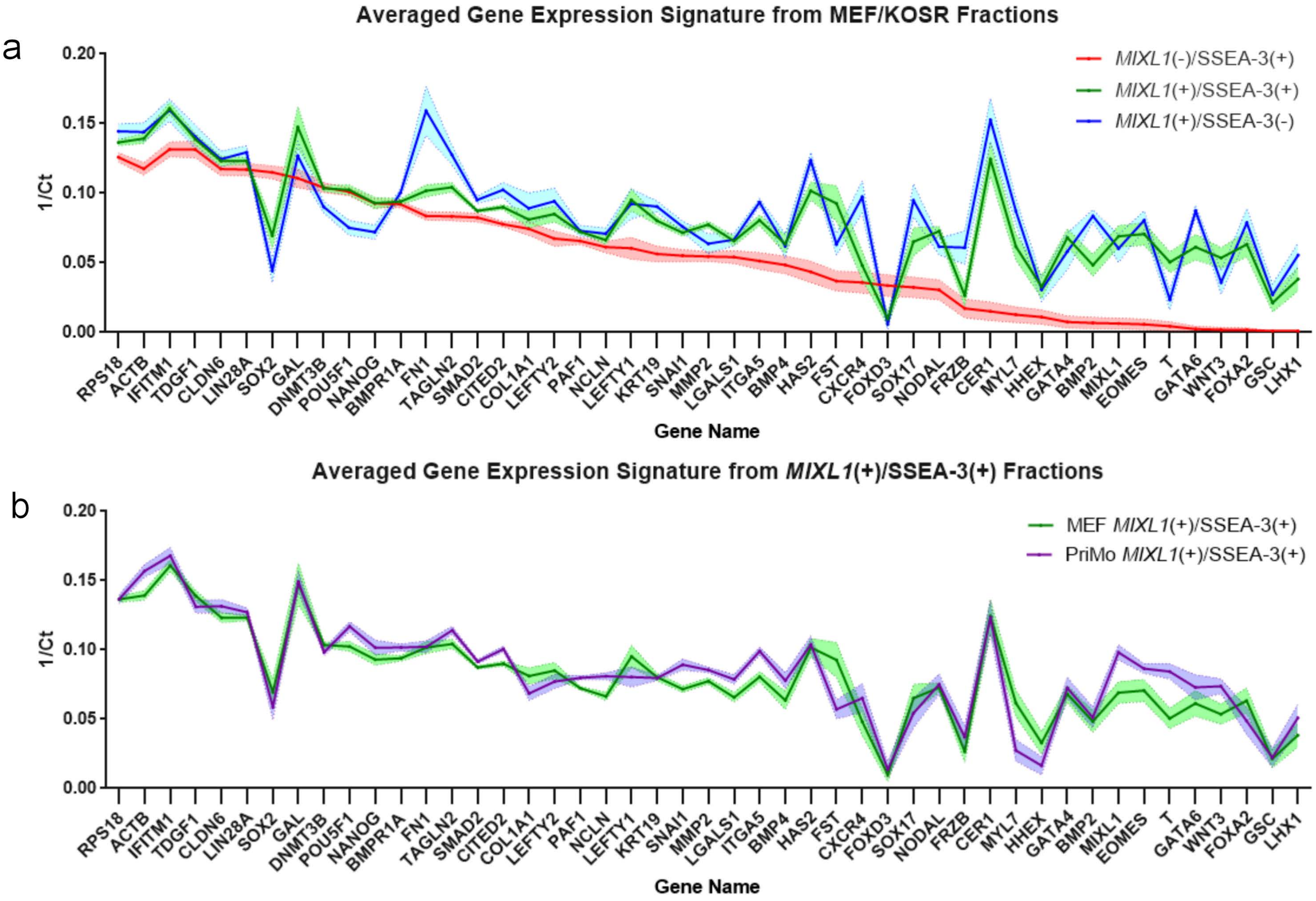
Averaged Gene Expression Signature from MEF/KOSR Fractions
a
0.20
0.15
0.10
0.05
0.00
1/Ct
MIXL1(-)/SSEA-3(+)
MIXL1(+)/SSEA-3(+)
MIXL1(+)/SSEA-3(-)
RPS18 ACTB IFITM1 TDGF1 CLDN6 LIN28A SOX2 GAL DNMT3B POU5F1 NANOG BMPR1A FN1 TAGLN2 SMAD2 CITED2 COL1A1 LEFTY2 PAF1 NCLN LEFTY1 KRT19 SNAI1 MMP2 LGALS1 ITGA5 BMP4 HAS2 FST CXCR4 FOXD3 SOX17 NODAL FRZB CER1 MYL7 HHEX GATA4 BMP2 MIXL1 EOMES T GATA6 WNT3 FOXA2 GSC LHX1
Gene Name
Averaged Gene Expression Signature from MIXL1(+)/SSEA-3(+) Fractions
b
0.20
0.15
0.10
0.05
0.00
1/Ct
MEF MIXL1(+)/SSEA-3(+)
PriMo MIXL1(+)/SSEA-3(+)
RPS18 ACTB IFITM1 TDGF1 CLDN6 LIN28A SOX2 GAL DNMT3B POU5F1 NANOG BMPR1A FN1 TAGLN2 SMAD2 CITED2 COL1A1 LEFTY2 PAF1 NCLN LEFTY1 KRT19 SNAI1 MMP2 LGALS1 ITGA5 BMP4 HAS2 FST CXCR4 FOXD3 SOX17 NODAL FRZB CER1 MYL7 HHEX GATA4 BMP2 MIXL1 EOMES T GATA6 WNT3 FOXA2 GSC LHX1
Gene Name

S4. a

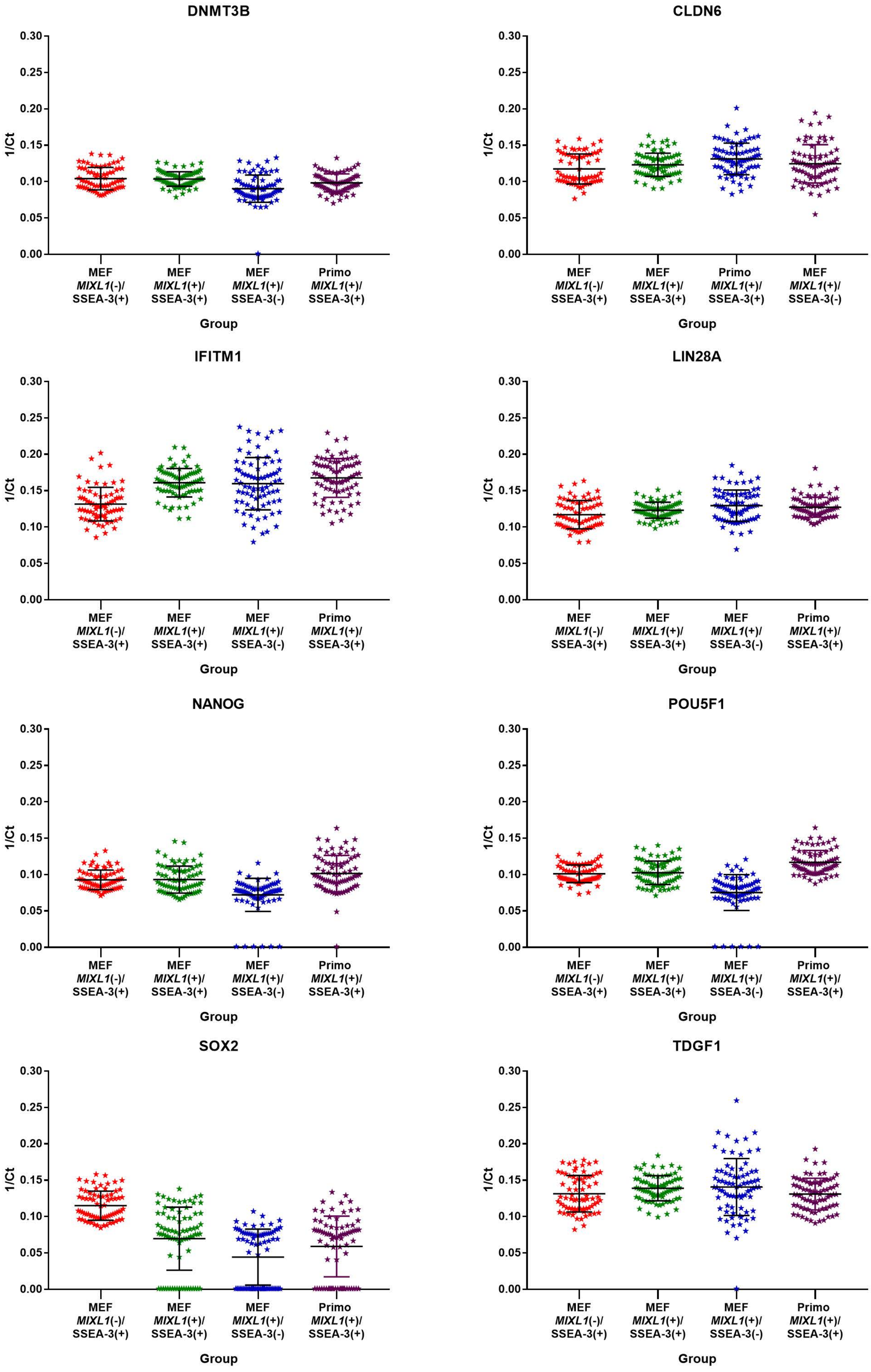
DNMT3B
CLDN6
IFITM1
LIN28A
NANOG
POU5F1
SOX2
TDGF1
1/Ct
0.00
0.05
0.10
0.15
0.20
0.25
0.30
MEF MIXL1(-)/ SSEA-3(+)
MEF MIXL1(+)/ SSEA-3(+)
MEF MIXL1(+)/ SSEA-3(-)
Primo MIXL1(+)/ SSEA-3(+)
Group

S4. b

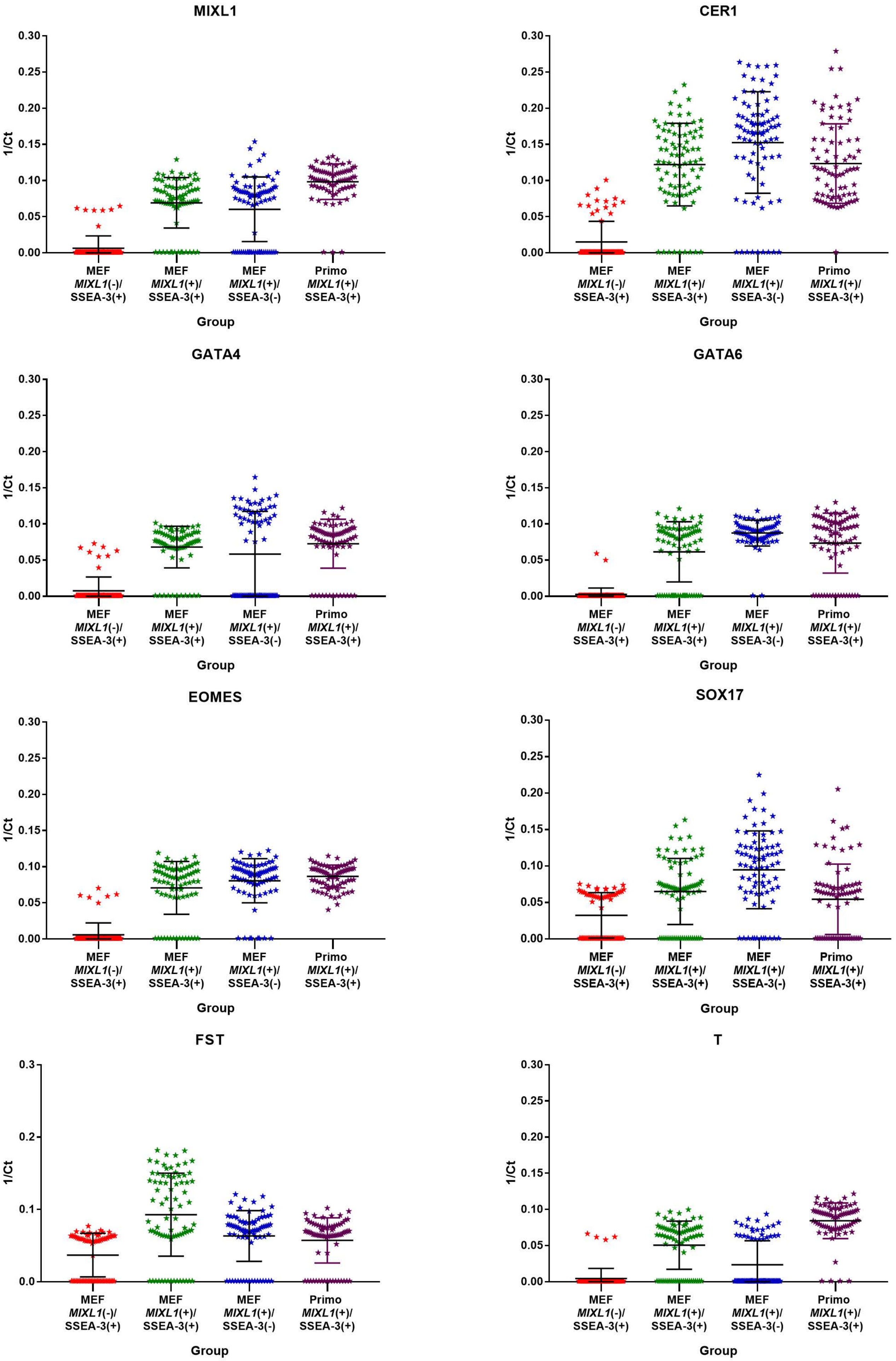
MIXL1
CER1
GATA4
GATA6
EOMES
SOX17
FST
T
1/Ct
0.00
0.05
0.10
0.15
0.20
0.25
0.30
0.0
0.1
0.2
0.3
MEF
MIXL1(-)/
SSEA-3(+)
MEF
MIXL1(+)/
SSEA-3(+)
MEF
MIXL1(+)/
SSEA-3(-)
Primo
MIXL1(+)/
SSEA-3(+)
Group

S4. c

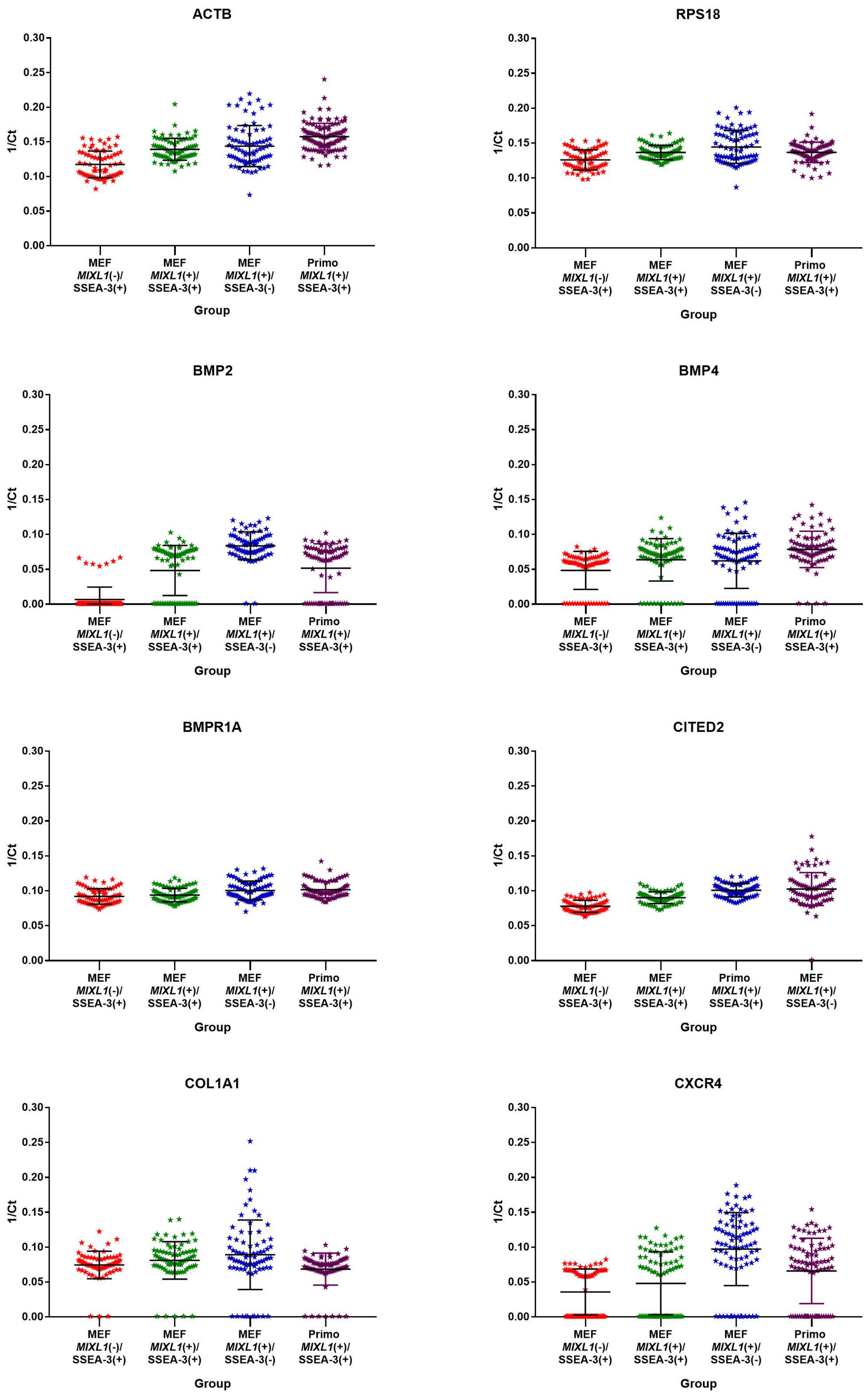

ACTB
RPS18
BMP2
BMP4
BMPR1A
CITED2
COL1A1
CXCR4
1/Ct
0.00
0.05
0.10
0.15
0.20
0.25
0.30
MEF MIXL1(-)/ SSEA-3(+)
MEF MIXL1(+)/ SSEA-3(+)
MEF MIXL1(+)/ SSEA-3(-)
Primo MIXL1(+)/ SSEA-3(+)
Group

S4. d

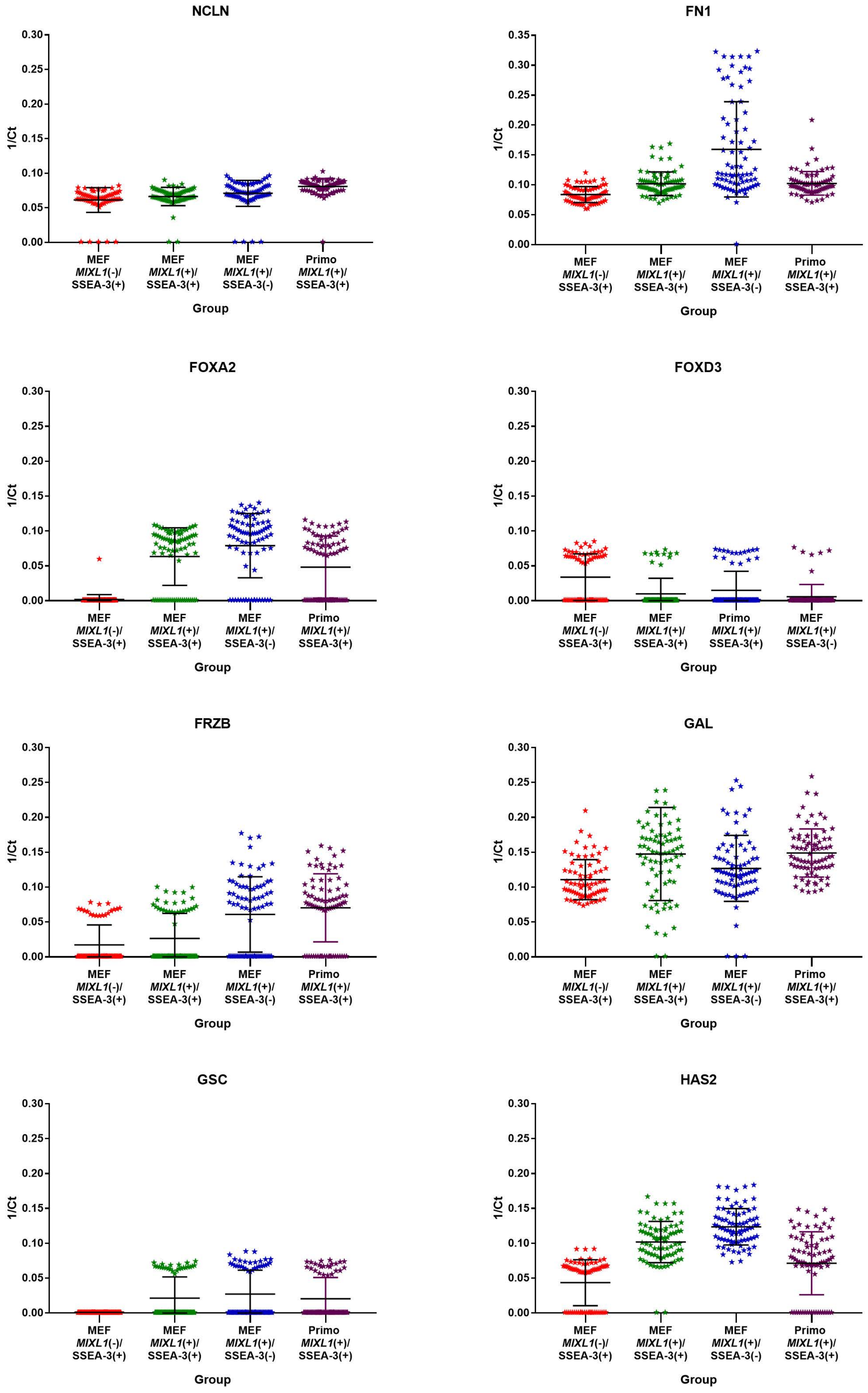
NCLN
1/Ct
0.30
0.25
0.20
0.15
0.10
0.05
0.00
MEF MIXL1(-)/ SSEA-3(+)
MEF MIXL1(+)/ SSEA-3(+)
MEF MIXL1(+)/ SSEA-3(-)
Primo MIXL1(+)/ SSEA-3(+)
Group
FN1
1/Ct
0.35
0.30
0.25
0.20
0.15
0.10
0.05
0.00
MEF MIXL1(-)/ SSEA-3(+)
MEF MIXL1(+)/ SSEA-3(+)
MEF MIXL1(+)/ SSEA-3(-)
Primo MIXL1(+)/ SSEA-3(+)
Group
FOXA2
1/Ct
MEF MIXL1(-)/ SSEA-3(+)
MEF MIXL1(+)/ SSEA-3(+)
MEF MIXL1(+)/ SSEA-3(-)
Primo MIXL1(+)/ SSEA-3(+)
Group
FOXD3
1/Ct
MEF MIXL1(-)/ SSEA-3(+)
MEF MIXL1(+)/ SSEA-3(+)
Primo MIXL1(+)/ SSEA-3(+)
MEF MIXL1(+)/ SSEA-3(-)
Group
FRZB
1/Ct
MEF MIXL1(-)/ SSEA-3(+)
MEF MIXL1(+)/ SSEA-3(+)
MEF MIXL1(+)/ SSEA-3(-)
Primo MIXL1(+)/ SSEA-3(+)
Group
GAL
1/Ct
MEF MIXL1(-)/ SSEA-3(+)
MEF MIXL1(+)/ SSEA-3(+)
MEF MIXL1(+)/ SSEA-3(-)
Primo MIXL1(+)/ SSEA-3(+)
Group
GSC
1/Ct
MEF MIXL1(-)/ SSEA-3(+)
MEF MIXL1(+)/ SSEA-3(+)
MEF MIXL1(+)/ SSEA-3(-)
Primo MIXL1(+)/ SSEA-3(+)
Group
HAS2
1/Ct
MEF MIXL1(-)/ SSEA-3(+)
MEF MIXL1(+)/ SSEA-3(+)
MEF MIXL1(+)/ SSEA-3(-)
Primo MIXL1(+)/ SSEA-3(+)
Group

S4. e

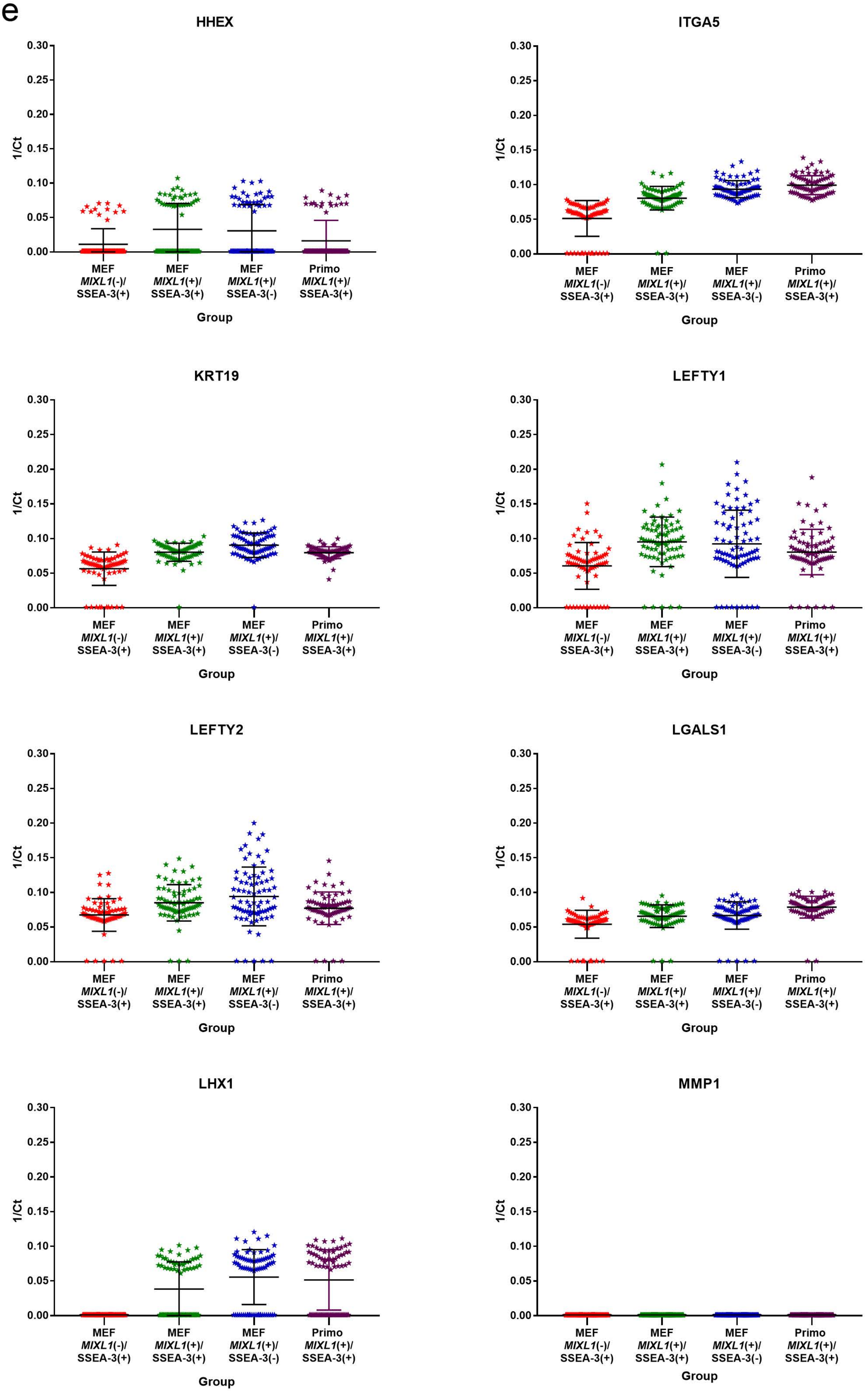
HHEX
ITGA5
KRT19
LEFTY1
LEFTY2
LGALS1
LHX1
MMP1
1/Ct
0.30
0.25
0.20
0.15
0.10
0.05
0.00
MEF *MIXL1*(-)/ SSEA-3(+)
MEF *MIXL1*(+)/ SSEA-3(+)
MEF *MIXL1*(+)/ SSEA-3(-)
Primo *MIXL1*(+)/ SSEA-3(+)
Group

S4. f

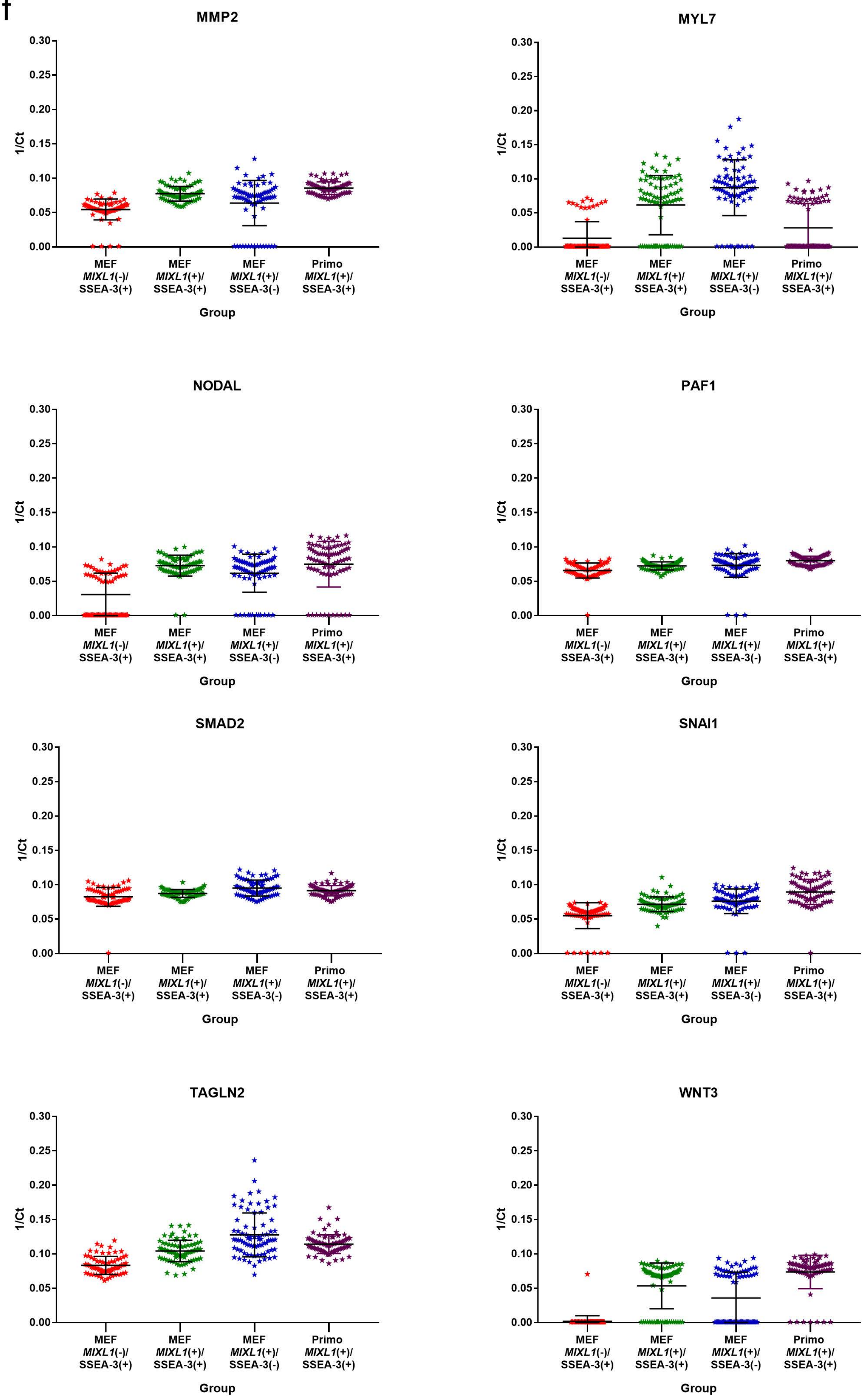
MMP2
MYL7
NODAL
PAF1
SMAD2
SNAI1
TAGLN2
WNT3
1/Ct
0.30
0.25
0.20
0.15
0.10
0.05
0.00
MEF MIXL1(-)/ SSEA-3(+)
MEF MIXL1(+)/ SSEA-3(+)
MEF MIXL1(+)/ SSEA-3(-)
Primo MIXL1(+)/ SSEA-3(+)
Group

a

S5

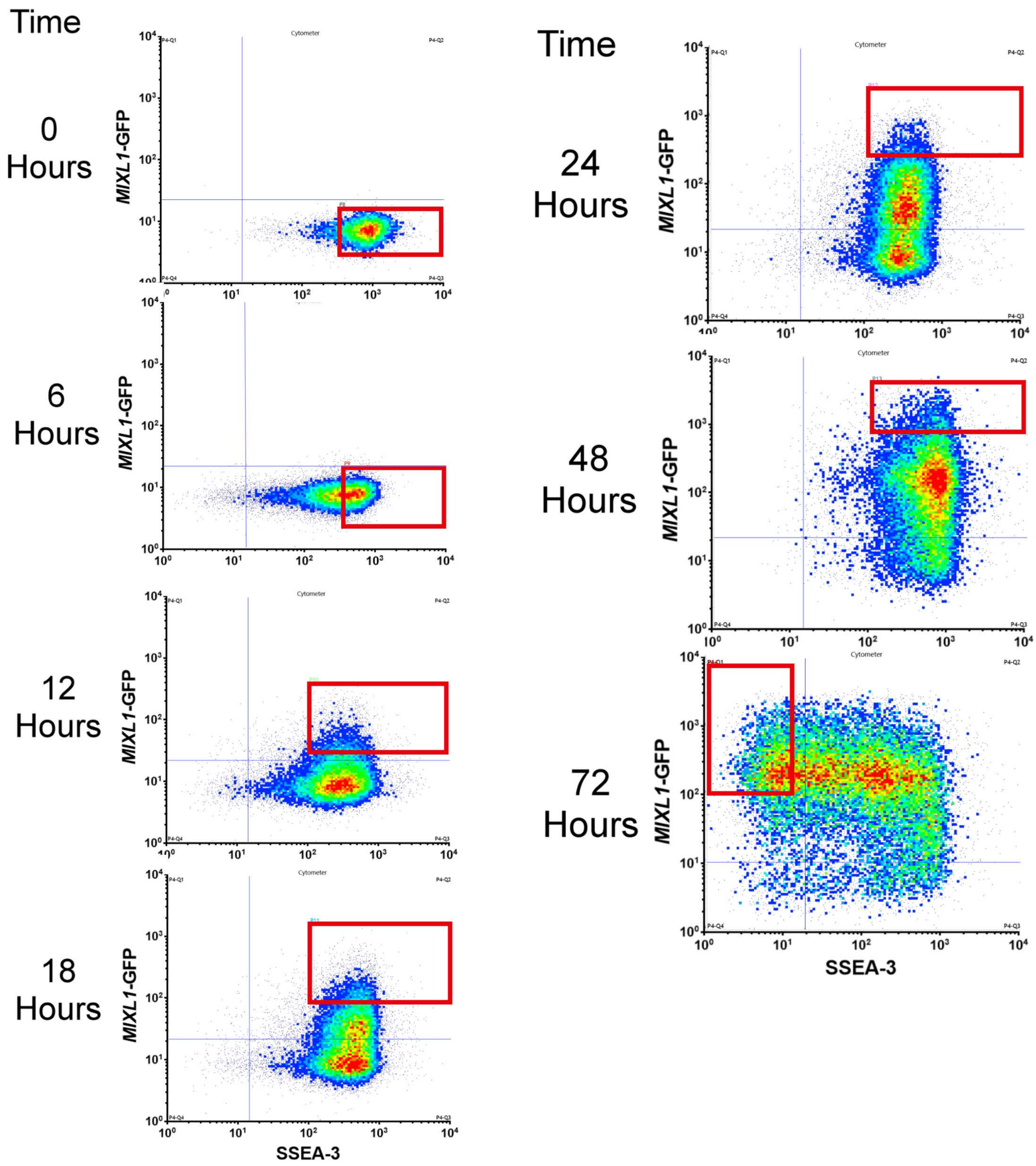

Time
0 Hours
6 Hours
12 Hours
18 Hours
Time
24 Hours
48 Hours
72 Hours
MIXL1-GFP
SSEA-3
Cytometer

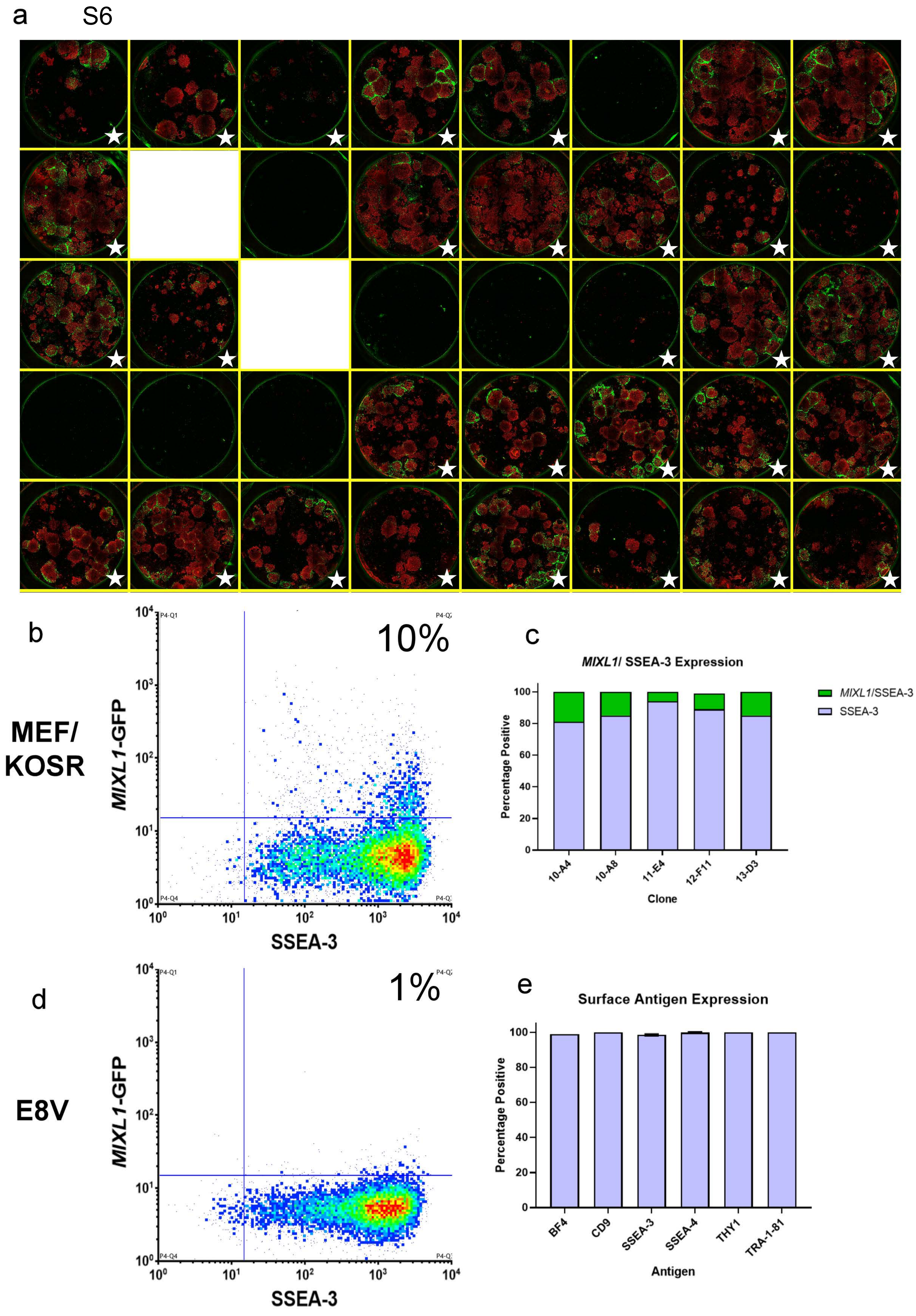
a
S6
b
MEF/ KOSR
10%
MIXL1-GFP
SSEA-3
c
MIXL1/ SSEA-3 Expression
Percentage Positive
MIXL1/SSEA-3
SSEA-3
10-A4
10-A8
11-E4
12-F11
13-D3
Clone
d
E8V
1%
MIXL1-GFP
SSEA-3
e
Surface Antigen Expression
Percentage Positive
BF4
CD9
SSEA-3
SSEA-4
THY1
TRA-1-81
Antigen

S7

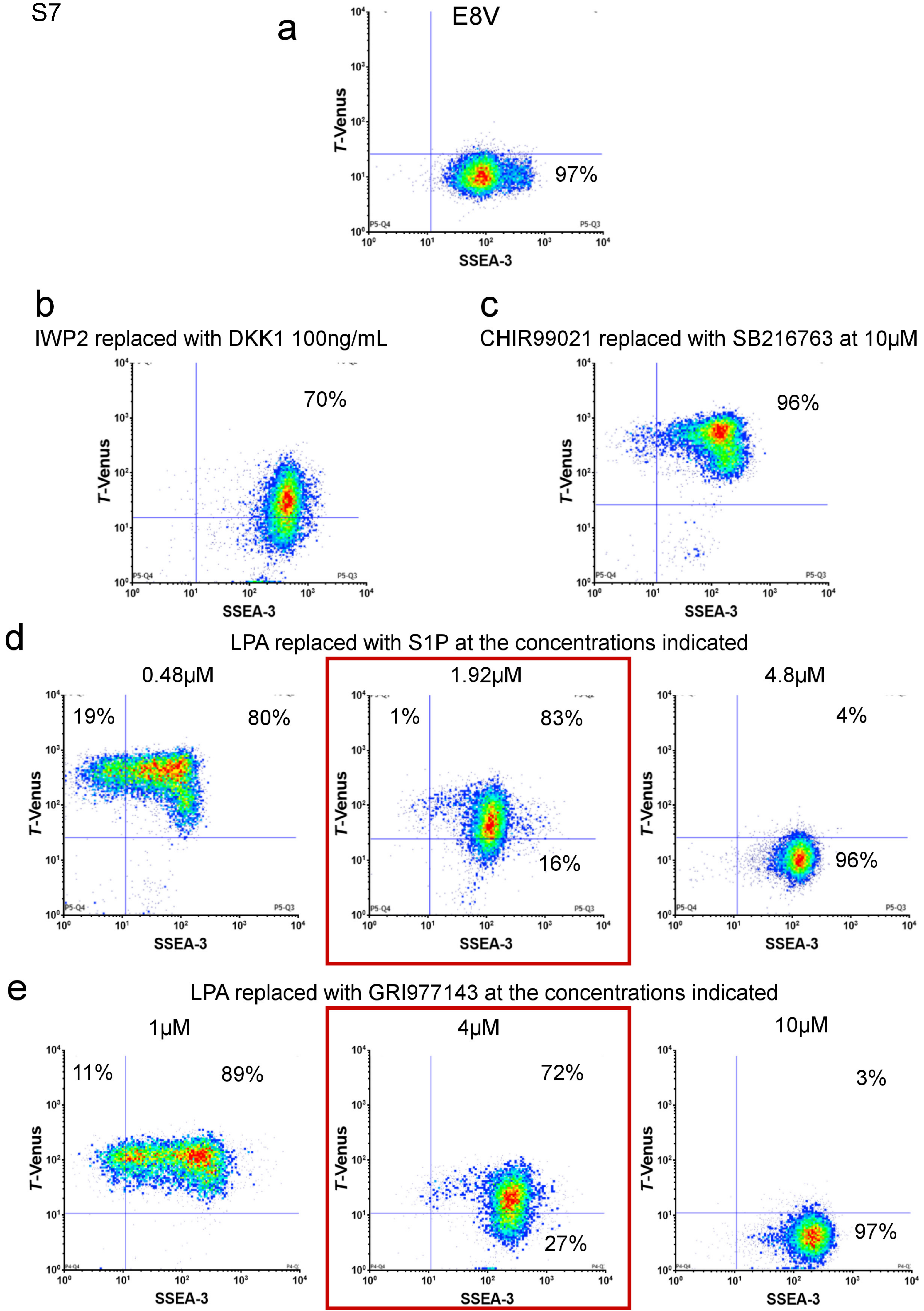

a
E8V
97%
T-Venus
SSEA-3
b
IWP2 replaced with DKK1 100ng/mL
70%
c
CHIR99021 replaced with SB216763 at 10μM
96%
d
LPA replaced with S1P at the concentrations indicated
0.48μM
19%
80%
1.92μM
1%
83%
16%
4.8μM
4%
96%
e
LPA replaced with GRI977143 at the concentrations indicated
1μM
11%
89%
4μM
72%
27%
10μM
3%
97%

S8

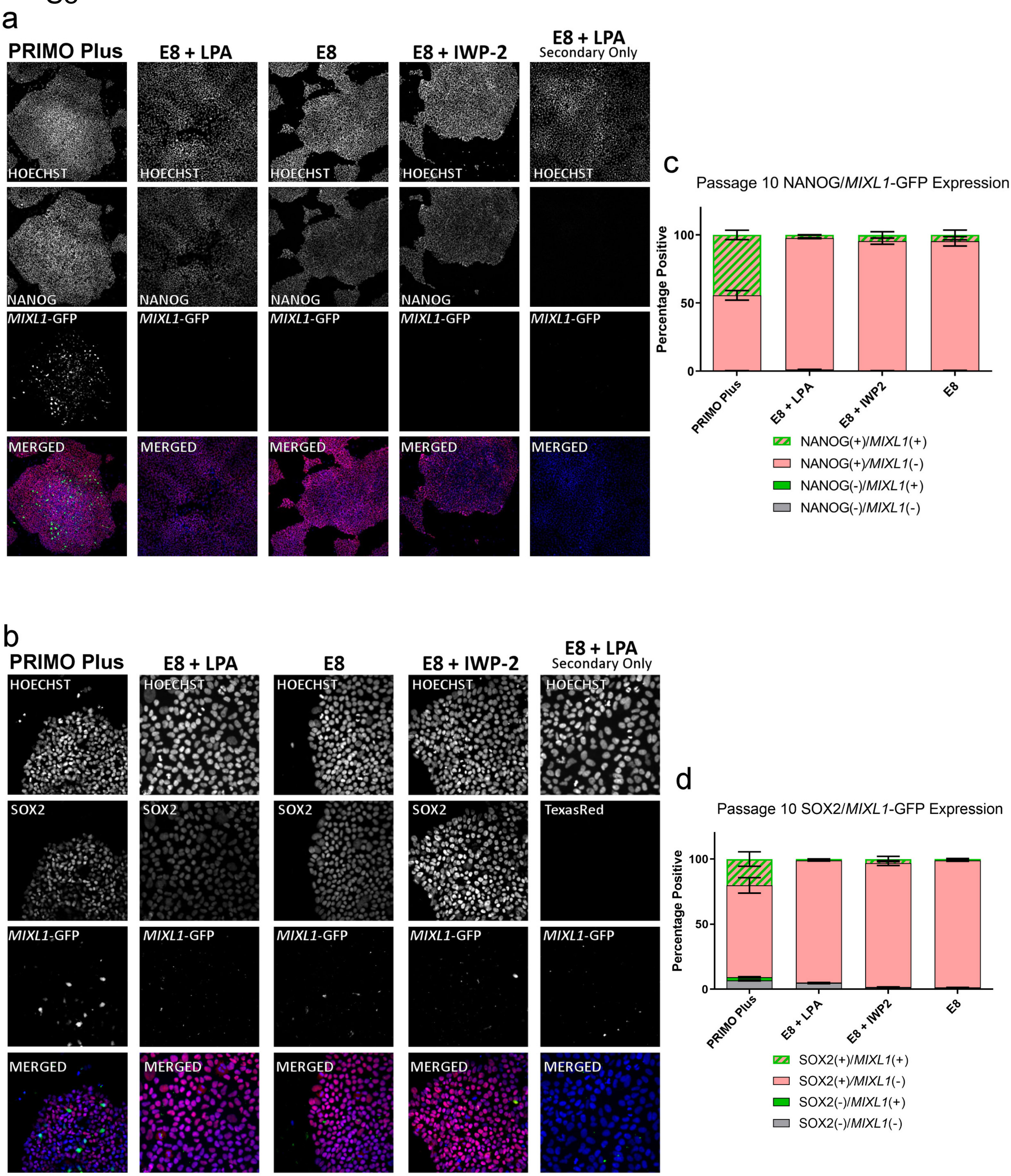
a
PRIMO Plus
E8 + LPA
E8
E8 + IWP-2
E8 + LPA
Secondary Only
HOECHST
NANOG
MIXL1-GFP
MERGED
b
SOX2
TexasRed
c
Passage 10 NANOG/MIXL1-GFP Expression
Percentage Positive
100
50
0
PRIMO Plus
E8 + LPA
E8 + IWP2
E8
NANOG(+)/MIXL1(+)
NANOG(+)/MIXL1(-)
NANOG(-)/MIXL1(+)
NANOG(-)/MIXL1(-)
d
Passage 10 SOX2/MIXL1-GFP Expression
SOX2(+)/MIXL1(+)
SOX2(+)/MIXL1(-)
SOX2(-)/MIXL1(+)
SOX2(-)/MIXL1(-)

### Table 1 Single Cell qPCR assay list

| Assay ID | Gene Symbol | RefSeq | Amplicon Length | Detects gDNA | Best Coverage |
|---|---|---|---|---|---|
| **Hs01060665_g1** | *ACTB* | NM_001101.3 | 63 | Yes | Yes |
| **Hs00154192_m1** | *BMP2* | NM_001200.2 | 60 | No | Yes |
| **Hs03676628_s1** | *BMP4* | NM_130850.2;NM_001202.3;NM_130851.2 | 116 | Yes | Yes |
| **Hs01034913_g1** | *BMPR1A* | NM_004329.2 | 94 | Yes | Yes |
| **Hs00193796_m1** | *CER1* | NM_005454.2 | 92 | No | Yes |
| **Hs01897804_s1** | *CITED2* | NM_001168388.2;NM_001168389.2;NM_006079.4 | 106 | Yes | Yes |
| **Hs00607528_s1** | *CLDN6* | NM_021195.4 | 154 | Yes | Yes |
| **Hs00164004_m1** | *COL1A1* | NM_000088.3 | 66 | No | Yes |
| **Hs00976734_m1** | *CXCR4* | NM_003467.2;NM_001008540.1 | 153 | No | No |
| **Hs00171876_m1** | *DNMT3B* | NM_001207055.1;NM_001207056.1;NM_175848.1;NM_175849.1;NM_175850.2;NM_006892.3 | 55 | No | Yes |
| **Hs00172872_m1** | *EOMES* | NM_001278183.1;NM_001278182.1;NM_005442.3 | 81 | No | Yes |
| **Hs01549976_m1** | *FN1* | NM_212482.1;NM_054034.2;NM_002026.2;NM_212478.1;NM_212474.1;NM_212476.1 | 81 | No | Yes |
| **Hs00232764_m1** | *FOXA2* | NM_021784.4;NM_153675.2 | 66 | No | Yes |
| **Hs00255287_s1** | *FOXD3* | NM_012183.2 | 78 | Yes | Yes |
| **Hs00173503_m1** | *FRZB* | NM_001463.3 | 108 | No | Yes |
| **Hs00246256_m1** | *FST* | NM_006350.3;NM_013409.2 | 108 | No | No |
| **Hs00544355_m1** | *GAL* | NM_015973.3 | 125 | No | Yes |
| **Hs00171403_m1** | *GATA4* | NM_002052.3 | 68 | No | Yes |
| **Hs00232018_m1** | *GATA6* | NM_005257.4 | 91 | No | Yes |
| **Hs00906630_g1** | *GSC* | NM_173849.2 | 100 | No | No |
| **Hs00193435_m1** | *HAS2* | NM_005328.2 | 63 | No | Yes |
| **Hs00242160_m1** | *HHEX* | NM_002729.4 | 110 | Yes | Yes |
| **Hs00705137_s1** | *IFITM1* | NM_003641.3 | 93 | Yes | Yes |
| **Hs01547673_m1** | *ITGA5* | NM_002205.2 | 54 | No | Yes |
| **Hs00761767_s1** | *KRT19* | NM_002276.4 | 116 | Yes | Yes |
| **Hs00764128_s1** | *LEFTY1* | NM_020997.3 | 136 | Yes | Yes |
| **Hs00745761_s1** | *LEFTY2* | NM_001172425.1;NM_003240.3 | 102 | Yes | Yes |
| **Hs00355202_m1** | *LGALS1* | NM_002305.3 | 63 | No | Yes |
| **Hs00232144_m1** | *LHX1* | NM_005568.3 | 60 | No | Yes |
| **Hs00702808_s1** | *LIN28A* | NM_024674.4 | 143 | Yes | Yes |
| **Hs00430824_g1** | *MIXL1* | NM_031944.1 | 152 | No | No |
| **Hs00899658_m1** | *MMP1* | NM_001145938.1;NM_002421.3 | 64 | No | Yes |

| Assay ID | Gene Symbol | RefSeq | Amplicon Length | Detects gDNA | Best Coverage |
|---|---|---|---|---|---|
| **Hs01548727_m1** | *MMP2* | NM_004530.4;NM_001127891.1 | 65 | No | Yes |
| **Hs01085598_g1** | *MYL7* | NM_021223.2 | 74 | No | Yes |
| **Hs04399610_g1** | *NANOG* | NM_024865.2 | 101 | Yes | No |
| **Hs00378379_m1** | *NCLN* | NM_020170.3 | 65 | No | Yes |
| **Hs00415443_m1** | *NODAL* | NM_018055.4 | 68 | No | Yes |
| **Hs00219496_m1** | *PAF1* | NM_019088.3;NM_001256826.1 | 100 | No | Yes |
| **Hs04260367_gH** | *POU5F1* | NM_001173531.1;NM_002701.4;NM_203289.4 | 77 | Yes | Yes |
| **Hs01375212_g1** | *RPS18* | NM_022551.2 | 93 | Yes | Yes |
| **Hs00183425_m1** | *SMAD2* | NM_001135937.2;NM_001003652.3;NM_005901.5 | 129 | No | No |
| **Hs00195591_m1** | *SNAI1* | NM_005985.3 | 66 | Yes | Yes |
| **Hs00751752_s1** | *SOX17* | NM_022454.3 | 149 | Yes | Yes |
| **Hs01053049_s1** | *SOX2* | NM_003106.3 | 91 | Yes | Yes |
| **Hs00610080_m1** | *T* | NM_001270484.1;NM_003181.3 | 132 | No | Yes |
| **Hs00761239_s1** | *TAGLN2* | NM_001277224.1;NM_001277223.1;NM_003564.2 | 163 | No | Yes |
| **Hs02339499_g1** | *TDGF1* | NM_003212.3;NM_001174136.1 | 170 | Yes | No |
| **Hs00902257_m1** | *WNT3* | NM_030753.4 | 76 | No | Yes |
